# Cryo-electron tomography of large biological specimens vitrified by plunge freezing

**DOI:** 10.1101/2021.04.14.437159

**Authors:** Felix J.B. Bäuerlein, Max Renner, Dana El Chami, Stephan E. Lehnart, José C. Pastor-Pareja, Rubén Fernández-Busnadiego

**Affiliations:** Department of Molecular Structural Biology, Max Planck Institute of Biochemistry, 82152 Martinsried, Germany; Institute of Neuropathology, University Medical Center Göttingen, 37099 Göttingen, Germany; Cluster of Excellence “Multiscale Bioimaging: from Molecular Machines to Networks of Excitable Cells” (MBExC), University of Göttingen, Germany; Collaborative Research Center SFB1190 “Compartmental Gates and Contact Sites in Cells”, University of Göttingen, 37073 Göttingen, Germany; Department of Chemistry, Umeå University, 901 87 Umeå, Sweden; Cellular Biophysics and Translational Cardiology Section, Heart Research Center Göttingen, University Medical Center Göttingen, 37075 Göttingen, Germany; Department of Cardiology and Pneumology, University Medical Center Göttingen, 37075 Göttingen, Germany; School of Life Sciences, Tsinghua University, 100084 Beijing, China; Tsinghua-Peking Center for Life Sciences, Tsinghua University, 100084 Beijing, China; Institute of Neurosciences, Consejo Superior de Investigaciones Científicas-Universidad Miguel Hernández, 03550 San Juan de Alicante, Spain; Faculty of Physics, University of Göttingen, 37077 Göttingen, Germany

## Abstract

Cryo-focused ion beam (cryo-FIB) milling allows thinning vitrified cells for high resolution imaging by cryo-electron tomography (cryo-ET). However, it remains challenging to apply this workflow to voluminous biological specimens such as tissues or particularly large mammalian cells, which usually require high-pressure freezing for vitrification. Here we show that adult mouse cardiomyocytes and dissected *Drosophila* tissues can be directly vitrified by plunge freezing upon a short incubation in 10% glycerol. This expedites subsequent cryo-FIB/ET, enabling systematic analyses of the molecular architecture of complex native specimens. Our data provides unanticipated insights into the molecular architecture of samples hitherto unexplored by cryo-ET.

## Introduction

Cryo-ET allows imaging the cell interior at molecular resolution (Wagner et al., 2017, Beck and Baumeister, 2016). This relies on the pristine preservation of cellular ultrastructure by vitrification (Dubochet and Sartori Blanc, 2001), which requires sufficiently fast cooling to prevent the re-organization of water molecules into crystalline ice (Dubochet et al., 1988). Cryo-FIB milling is currently the method of choice to thin down vitrified cells into electron-transparent lamellae (Rigort et al., 2012a, Marko et al., 2007, Strunk et al., 2012, Hayles et al., 2010) to enable high-quality cryo-ET imaging.

To date, cryo-FIB/ET has been applied mainly to isolated cells in culture (Wagner et al., 2017, Beck and Baumeister, 2016), which can in principle be conveniently vitrified by plunge freezing into a cryogen (Dubochet et al., 1988). Plunge frozen cells are embedded in a thin film of buffer or medium, and can thus be directly thinned down by cryo-FIB. In practice, however, most cultured mammalian cells are not fully vitrified by plunge freezing, as their core regions are too thick (⪆ 10 μm) for efficient heat transfer (Mahamid et al., 2016). To achieve vitrification of specimens up to ∼200 μm-thick, it is possible to resort to high-pressure freezing (Studer et al., 2008). However, this results in specimens embedded in a thick block of ice that prevents direct cryo-FIB milling, requiring complex procedures such as cryo-sectioning or cryo-lift out (Mahamid et al., 2015, Schaffer et al., 2019, Al-Amoudi, 2004, Hsieh et al., 2014, De Winter et al., 2021, Hayles et al., 2010). Furthermore, since no surface topography is visible in high-pressure frozen samples, the identification of target structures requires correlative light-microscopy. Altogether, the throughput of current cryo-ET tissue workflows is insufficient to address complex biological questions.

To achieve full vitrification of cultured cells, we routinely incubate them in a solution of 10% glycerol prior to plunge freezing (Bäuerlein et al., 2017). Morphological subtomogram averaging analyses indicate that glycerol treatment is not detrimental to cellular architecture (Bäuerlein et al., 2017, Guo et al., 2018). In fact, glycerol is a widely used cryo-protectant, as it substantially increases the glass transition temperature (Morris et al., 2012) and has minimal influence on protein structure and cell function (Ye et al., 2006, Sousa, 1995). Glycerol is naturally produced by cells (Alves-Bezerra and Cohen, 2017, Zhang et al., 2018), and is commonly used in human medicine (Liu, 2011, Robergs and Griffin, 1998, Righetti et al., 2004). Furthermore, glycerol production is a well-documented adaptation to survive freezing winter temperatures in some insects and vertebrates (Zachariassen, 1985, Rexer-Huber et al., 2011). Here we show that a brief incubation in 10% glycerol is also sufficient to allow vitrification of large mammalian cells such as adult mouse cardiomyocytes, as well as native tissues dissected from third instar (L3) larvae of the fruit fly *Drosophila melanogaster*. Cryo-ET analysis of these hitherto unexplored systems paves the way to address a myriad biological questions at molecular resolution.

## Results

### Cryo-ET of native *Drosophila melanogaster* tissues

*Drosophila* is advantageous for cryo-FIB/ET because its organs are relatively small and can be conveniently dissected. First, we tested whether the *Drosophila* nervous system could be vitrified by plunge freezing. To that end, we dissected cephalic complexes, containing the central nervous system (CNS) and peripheral nerves. Isolated cephalic complexes were deposited on conventional carbon-coated EM grids and plunge frozen upon 2-5 min incubation in PBS with (Fig. 1g-l) or without 10% glycerol (Fig. 1a-f). Visual examination by cryo-FIB and cryo-scanning electron microscopy (cryo-SEM) revealed a high degree of anatomical detail (Fig. 1a, b, g, h), allowing direct identification of structures of interest. Initially we targeted the peripheral nerves, recognizable as long tubules projecting from the ventral nerve cord (VNC; Fig. 1a, b, g, h). The nerves were ∼15 μm in diameter, and thus comparable in thickness to the perinuclear region of cultured mammalian cells. We prepared cryo-FIB lamellae of these regions using similar milling parameters as with cultured cells (Bäuerlein et al., 2017) (Fig. 1c, d, i, j), and inspected them by cryo-transmission electron microscopy (cryo-TEM) imaging and diffraction analyses. Low magnification imaging of lamellae produced in the absence of glycerol revealed high-contrast Bragg reflections indicative of ice crystal formation (Fig. 1e). These features were striking in tomographic tilt series (Supplementary Video 1). Consistently, diffraction analysis of these lamellae showed spots and sharp rings characteristic of crystalline ice (Dubochet et al., 1988) (Fig. 1f). In contrast, lamellae produced from nerves incubated with glycerol had a smooth appearance at low and high magnifications (Fig. 1k, Supplementary Video 1). Furthermore, diffraction analysis (Fig. 1l) revealed diffuse rings characteristic of vitreous ice in these samples (Dubochet et al., 1988, Blackman and Lisgarten, 1997). Therefore, the cooling rates achieved by plunge freezing were sufficient to vitrify thin peripheral nerves with the aid of glycerol.

**Fig. 1:**
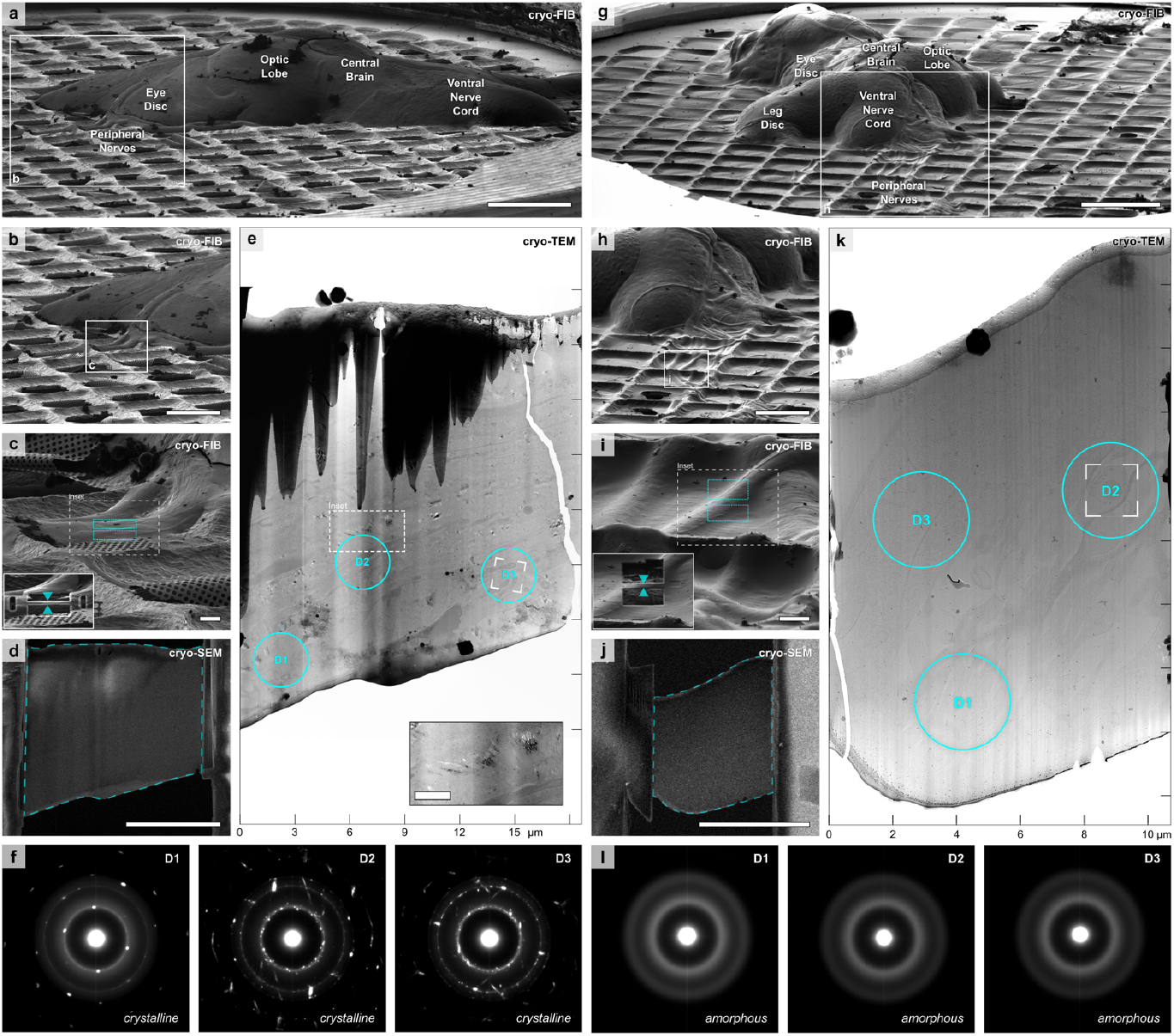
Short incubation in 10% glycerol allows vitrification of Drosophila peripheral nerves by plunge freezing. **(a)-(f)** Cryo-FIB lamella preparation and diffraction analysis of nervous tissue incubated 2-5 min in PBS only. **(a)** Cryo-FIB-induced secondary electron image of a cephalic complex (1.2×0.6×0.2 mm3), including the central nervous system, plunge frozen on an EM grid. **(b)** Magnification of the area boxed in (a) showing peripheral nerves emerging from the ventral nerve cord. **(c)** Magnification of the area boxed in (b). Blue rectangles indicate patterns for tissue removal by cryo-FIB milling. The inset shows the indicated region during lamella preparation (blue arrows). **(d)** Top view cryo-SEM image of the final lamella shown in (c). The blue line indicates the lamella area shown in (e). **(e)** Cryo-TEM overview of the final lamella shown in (c, d). Blue circles indicate the regions where vitrification was assessed by electron diffraction (f). The inset shows a magnification of the region boxed in the main panel with a white broken line. This region shows artifacts (black/white Bragg reflections) caused by diffraction of the electron beam in crystalline ice. The box within D3 marks the area where the tilt series shown in Supplementary Video 1 (left) was recorded. **(f)** Diffraction patterns recorded on the regions D1-D3 marked in (e), showing predominantly crystalline ice, even close to the lamella surface (D1). **(g)-(l)** Cryo-FIB lamella preparation and diffraction analysis of nervous tissue incubated 2-5 min in PBS with 10% glycerol. **(g)** Cryo-FIB-induced secondary electron image of a cephalic complex (1.0×0.6×0.2 mm3) plunge frozen on an EM grid. **(h)** Magnification of the area boxed in (g) showing peripheral nerves emerging from the ventral nerve cord. **(i)** Magnification of the area boxed in (h). Blue rectangles indicate patterns for tissue-removal by cryo-FIB milling. The inset shows the indicated region during lamella preparation (blue arrows). **(j)** Top view cryo-SEM image of the final lamella shown in (i). The blue line indicates the lamella area shown in (k). **(k)** Cryo-TEM overview of the final lamella shown in (i, j). Blue circles indicate regions where vitrification was assessed by electron-diffraction (l). The lamella appears smooth, without artifacts induced by diffraction of the electron beam in crystalline ice. The box within D2 marks the area where the tilt series shown in Supplementary Video 1 (right) was recorded. **(l)** The diffraction patterns of the regions D1-D3 marked in (k) show diffuse rings characteristic of vitreous ice throughout the complete lamella. Scale bars: 200 μm (a, g), 100 μm in (b, h), 10 μm (c, d, i, j), 1 μm (e, inset).

A common difficulty in cryo-FIB/ET is to visualize and subsequently target appropriate tomographic regions in low magnification lamella overviews. The contrast of these images is dominated by curtaining artifacts and thickness variations (Bäuerlein et al., 2017, Guo et al., 2018, Arnold et al., 2016) (Fig. 1e, k), arising from the different milling efficiency on different cellular components and the limited focus depth of the ion beam, respectively (Rigort et al., 2012a). Increasing image contrast makes these artifacts even more pronounced (Extended Data Fig. 1a, b). This is especially detrimental for analyzing cryo-FIB lamellae of tissues, given their rich biological complexity. To address this issue, we developed an algorithm termed LisC (“Lamella *in silico* Clearing”) to remove obscuring features from lamella overviews. First, a high-pass filter is applied to remove the low frequency brightness modulation caused by long-range thickness variations (Extended Data Fig. 1c, d). Then, a line filter is applied in Fourier space to remove line artifacts caused by curtaining (Extended Data Fig. 1e, f). This results as well in the formation of ringing artifacts near high-contrast features, such as ice crystal contamination or the vacuum background (Extended Data Fig. 1e). However, when these features are masked by a local gray value average (Extended Data Fig. 1h), ringing artifacts are minimized and Fourier line filtering results in high-contrast images where the structural features of the lamella can be clearly discerned (Extended Data Fig. 1g, Supplementary Video 2).

We next examined lamellae of vitrified glycerol-treated nerves by cryo-ET. LisC filtering of lamella overview images allowed selecting areas for cryo-ET data collection based on a detailed histological analysis (Fig. 2b, Extended Data Fig. 2e). Cryo-ET revealed an excellent preservation of cell morphology (Fig. 2c-d), consistent with our previous studies in glycerol-treated cultured cells (Bäuerlein et al., 2017, Guo et al., 2018). Importantly, tomograms of vitrified nerves showed molecular details of tissue-specific structures that cannot be reproduced in culture, such as the wrapping of axons by perineurial glia or the intricate architecture of the basement membrane (Fig. 2c, d, Supplementary Video 3). It was routinely possible to record multiple tomograms within one lamella, providing a broad overview of different regions of the tissue (Extended Data Fig. 2). These data indicate that peripheral nerves ∼15 μm in diameter can be vitrified by plunge freezing upon a brief incubation in glycerol, allowing direct cryo-ET analysis of native tissue architecture.

**Fig. 2:**
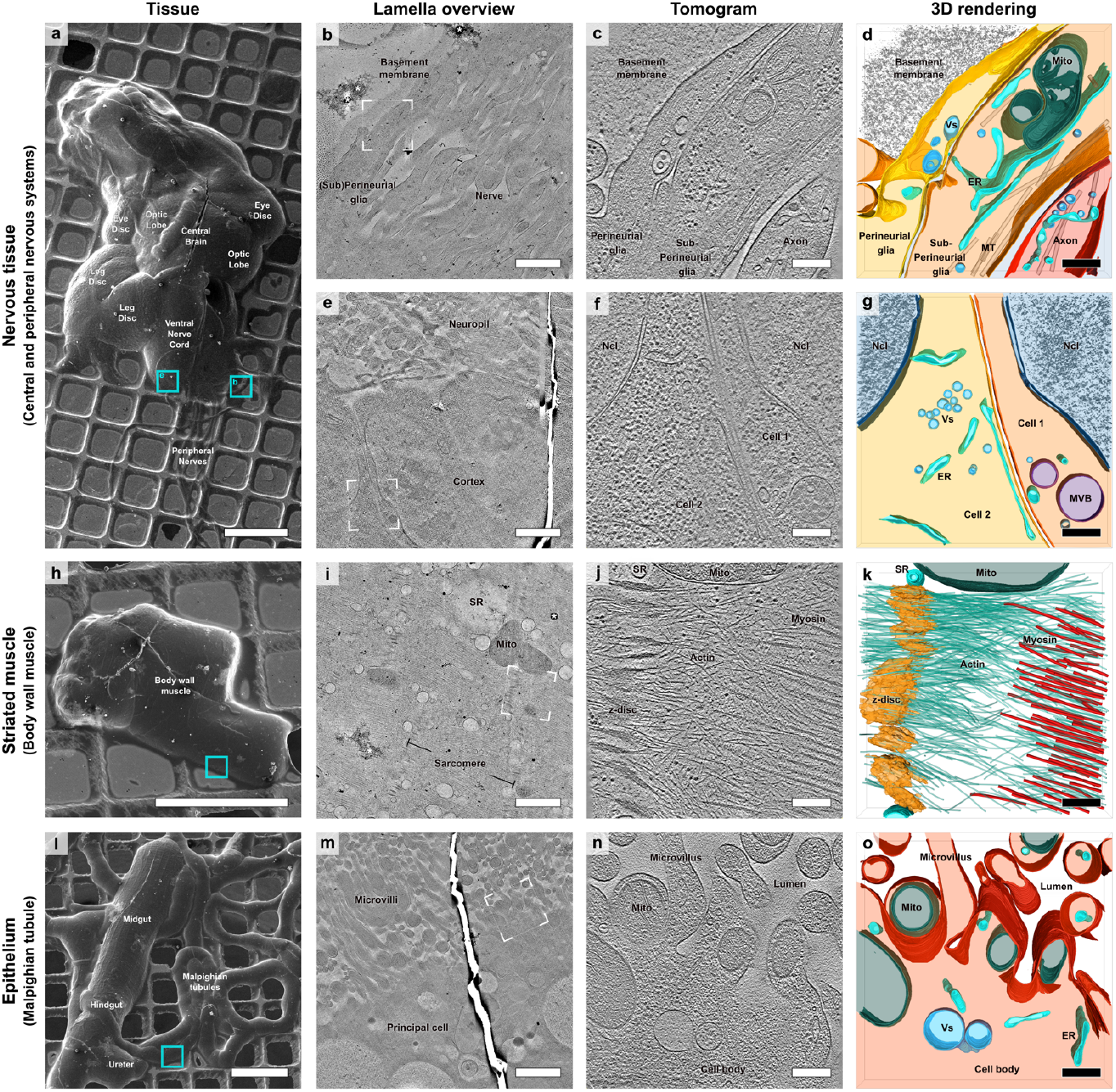
Gallery of Drosophila tissues analyzed by cryo-FIB/ET upon vitrification by plunge freezing. Rows represent different tissue types, and columns show different visualizations: cryo-SEM (first column), cryo-TEM lamella overviews filtered by the LisC algorithm (second column), cryo-ET slices (third column) and 3D renderings (fourth column). **(a)-(d)** Peripheral nervous system (peripheral nerve). **(a)** Top view cryo-SEM image of Drosophila cephalic complex shown in Fig. 1g. Anatomical features can be directly observed and are annotated. Blue boxes indicate regions of lamella preparation of the peripheral (b) and central (e) nervous system. **(b)** Cryo-TEM overview of a lamella prepared in the area marked in (a) on a peripheral nerve (see also Extended Data Fig. 2). The lamella contains multiple axons, together with sub-perineurial and perineurial glial cells and a basement membrane wrapping the nerve. * mark crystalline ice contamination deposited on top of the lamella. **(c)** 1.7 nm-thick slice of a tomogram recorded in the area marked in (b), showing an axon, the sub-perineurial and perineurial glia and the basement membrane. **(d)** 3D rendering of the tomogram in (c). Warm colors represent different cells, cold colors represent subcellular structures: axon (red), sub-perineurial glia (orange), perineurial glia (yellow), basement membrane (gray), interstitial space (transparent light blue), endoplasmic reticulum (ER, turquoise), mitochondria (Mito, dark green), microtubules (MT, transparent gray), vesicles (Vs, light blue). See also Supplementary Video 3. **(a, e-g)** Central nervous system (VNC cortex). **(e)** Cryo-TEM overview of a lamella prepared in the area marked in (a). Cell bodies can be seen in the bottom part of the image, while the neuropil is visible on the top. **(f)** 1.7 nm-thick slice of a tomogram recorded in the area marked in (e) showing two adjacent cells of the ventral nerve cord. Ncl: nucleus. **(g)** 3D rendering of the tomogram shown in (f). Cell 1 (orange), cell 2 (yellow), nucleus (Ncl, dark blue), endoplasmic reticulum (ER, turquoise), multi vesicular bodies (MVB, violet), vesicles (Vs, light blue). See also Supplementary Video 4. **(h)-(k)** Striated muscle (body wall muscle). **(h)** Top view cryo-SEM image of a body wall muscle from an abdominal segment. A blue box indicates the region targeted for lamella preparation. **(i)** Cryo-TEM overview of a lamella prepared in the area marked in (h). The lamella shows the sarcomeres, the sarcoplasmic reticulum (SR) and mitochondria (Mito). **(j)** 1.7 nm-thick slice of a tomogram recorded in the area marked in (i), showing the interplay of myosin and actin filaments associated to the z-disc. **(k)** 3D rendering of the tomogram in (j). Myosin fibrils (red), actin fibrils (green), z-disc (yellow), sarcoplasmic reticulum (SR, turquoise), mitochondria (Mito, dark green). See also Supplementary Video 5. (l)-(o) Epithelium (Malpighian tubule). **(l)** Top view cryo-SEM image of the Malpighian tubules, associated via the ureter to the gut. A blue box indicates the region targeted for lamella preparation. **(m)** Cryo-TEM overview of a lamella prepared in the area marked in (l). The lamella shows the luminal side of a principal cell body of the Malpighian tubule, with the cell body in the lower part of the image and the microvilli ranging into the tubule lumen in the top (see also Extended Data Fig. 3). **(n)** 1.7 nm-thick slice of a tomogram recorded in the area marked in (m), showing the luminal side of the principal cell. Mito: mitochondrion. **(o)** 3D rendering of the tomogram shown in (n). Principal cell (red), endoplasmic reticulum (ER, turquoise), mitochondria (Mito, dark green), vesicles (Vs, light blue). See also Supplementary Video 6. Scale bars: 200 μm (a, h, l), 1.5 μm (b, e, i, m), 250 nm (c, d, f, g, j, k, n, o).

However, other *Drosophila* tissues are substantially thicker than peripheral nerves, which may complicate vitrification and cryo-FIB milling. Therefore, we investigated whether superficial regions of the CNS (∼200 μm thick) could also be vitrified by plunge freezing in 10% glycerol. We prepared cryo-FIB lamellae of the VNC cortical region (Fig. 2a, e) using a 5-10 fold higher ion beam current to efficiently remove large amounts of material during the initial milling step. Cryo-ET imaging of the resulting lamellae revealed smooth membranes and high density of cytosolic macromolecules, with no signs of crystalline ice (Fig. 2e-g, Supplementary Video 4). Thus, glycerol incubation also allowed vitrification and cryo-ET imaging of superficial regions of the CNS.

To explore whether our approach was also applicable beyond nervous tissue, we examined body wall muscles. Dissected muscles with attached innervations were also directly amenable to direct cryo-ET imaging upon cryo-FIB milling (Fig. 2h-k). Low magnification images showed sarcomeres intercalated with mitochondria and the sarcoplasmic reticulum (Fig. 2i). Cryo-ET analysis of the sarcomere revealed the detailed organization of thick and thin filaments, as well as their connection to the Z-disk (Fig. 2j, k, Supplementary Video 5). Therefore, our approach allows imaging at molecular resolution complete neuromuscular circuits of *Drosophila* larvae, from the neuronal cell bodies in the VNC (Fig. 2a, e-g), their axonal projections forming the peripheral nerves (Fig. 2a-d) to the body wall muscles they innervate (Fig. 2h-k).

As an example of epithelial tissue, we investigated the Malpighian tubules, functional equivalent of the vertebrate kidney. Plunge-frozen dissected Malpighian tubules were directly milled by cryo-FIB (Fig. 2l-o). Lamellae overviews and cryo-tomograms showed the exquisite complexity of this polarized tissue, from the basal system of labyrinth channels to the apical microvilli (Fig. 2l-o, Extended Data Fig. 3, Supplementary Video 6). Occasionally, crystalline ice was observed in the tubule lumen, where the lower solute concentration provides lower cryo-protection, but not within the cell interior. Therefore, a short incubation in 10% glycerol allows vitrification of superficial areas of nervous, muscle and epithelial *Drosophila* tissues by plunge freezing, greatly facilitating subsequent cryo-FIB milling and cryo-ET imaging.

### Structural analysis of adult mouse cardiomyocytes

We next sought to explore the validity of our approach for large mammalian cells such as cardiomyocytes, the contractile units of the heart. Mouse cardiomyocytes are typically rod-shaped with dimensions of ∼20 μm in diameter and over 100 μm in length (Bensley et al., 2016, Oliver-Gelabert et al., 2020). Mature cardiomyocytes are morphologically unique, with a large fraction of their cellular volume occupied by highly-ordered sarcomeres interspersed by dense rows of mitochondria. Excitation-contraction coupling is mediated by an extensive network of transverse tubules (T-tubules), plasma membrane invaginations criss-crossing the cell (Wagner et al., 2012). T-tubules form membrane contact sites known as “junctional membrane complexes (JMCs)” with sarco/endoplasmic reticulum (SER) cisternae (Lehnart and Wehrens, 2022). At JMCs, Ca2+ influx through plasma membrane dihydropyridine receptors triggers a much larger SER Ca2+ release through type 2 ryanodine receptors (RyR2) to drive the contraction of the sarcomeric actomyosin machinery. Recent cryo-ET studies have analyzed native sarcomeric structure within neonatal rat cardiomyocytes (Burbaum et al., 2021) and homogenized skeletal muscle (Wang et al., 2021), but the molecular architecture of adult mature cardiomyocytes remains unexplored. In particular, the specific macromolecular arrangement of JMCs that enable Ca2+-induced Ca2+ release is poorly understood.

To shed light on this issue, we obtained intact living, relaxed ventricular cardiomyocytes from collagenase-digested adult mouse hearts, deposited them on EM grids and plunge-froze them upon one minute incubation in buffer containing 10% glycerol. Cardiomyocytes showed a characteristic rod-like morphology, with sharp-edges indicative of morphologically intact cells (Wagner et al., 2014) (Fig. 3a, b). Subsequent cryo-FIB milling produced large lamellae (Fig. 3c, d) showing a rich variety of cellular features and no visible signs of crystalline ice. Extensive alternating arrays of parallel myofibrils and mitochondria were intercalated by T-tubules of diverse morphology (Fig. 3d).

**Fig. 3:**
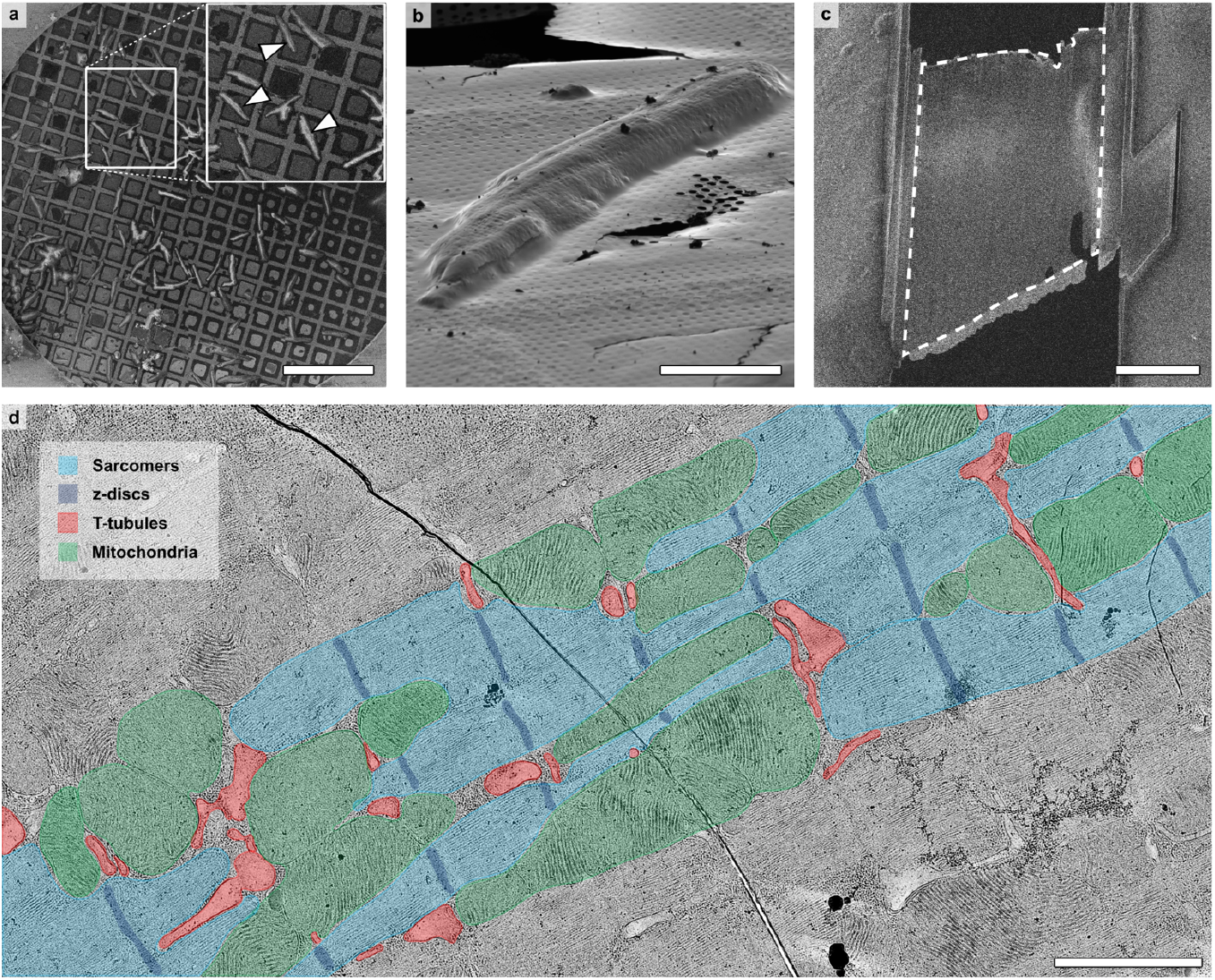
Preparation of plunge-frozen adult mouse cardiomyocytes for cryo-ET. **(a)** SEM overview of an EM grid containing plunge-frozen mouse ventricular cardiomyocytes (arrowheads). **(b)** FIB-induced image of a single cardiomyocyte. **(c)** SEM image of a fine-milled lamella of less than 100 nm thickness. **(d)** LisC-cleared low magnification TEM overview of a different lamella. A region of the lamella is pseudo-colored to indicate the most abundant cellular features, i.e. mitochondria (green), sarcomeres (light blue), z-discs (dark blue) and T-tubules (red). Scale bars: 500 μm (a), 25 μm (b), 10 μm (c), 2 μm (d).

Cryo-ET analysis of T-tubules revealed abundant electron-dense particles on their extracellular side (Fig. 4e-g, black arrowheads). T-tubules were often wrapped by SER cisternae (junctional SER or “jSER”) establishing JMCs, in many instances also tightly coupled with mitochondrial membranes on their off-T-tubule side (Fig. 4a, g). Interestingly, we observed two morphologically distinct domains at JMCs (Fig. 4a, b). A first domain (“narrow JMC”) showed an average distance between T-tubules and jSER membranes of ∼12 nm (Fig. 4c). Close inspection of these regions revealed numerous filamentous tethering densities linking the T-tubules and jSER membranes (Fig. 4e, white arrowheads). Other JMC regions, in some cases directly adjacent to narrow JMCs, showed an approximately double intermembrane distance (“wide JMC”; Fig. 4c). Intriguingly, the intermembrane space of wide JMCs showed densities parallel to the membranes, which corresponded in size to RyR2s (Fig. 4f, white arrows). In contrast, the ∼12 nm intermembrane distance of narrow JMCs is unlikely to accommodate RyR2s given their large size (Peng et al., 2016). Narrow and wide JMCs were not only distinguishable by T-tubule-jSER distance, but also by jSER morphology. jSER was extremely thin (∼8 nm wide) at narrow JMCs, leaving barely any luminal space (Fig. 4d, e). jSER was thicker at wide JMCs and showed an electron dense structure in the lumen reminiscent to the calsequestrin polymer layer (Fig. 4d, f, black arrows).

**Fig. 4:**
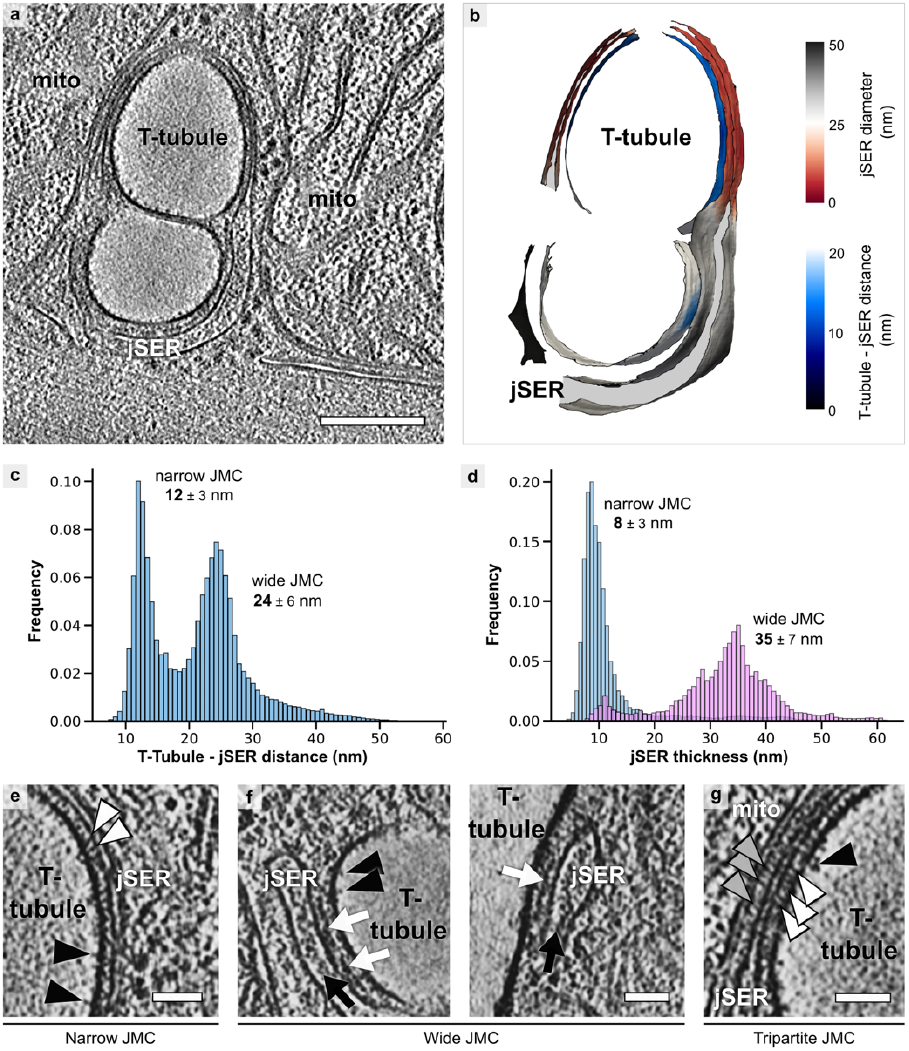
Structural analysis of junctional membrane complexes. **(a)** Tomographic slice of a JMC showing a double T-tubule recorded on a cryo-FIB lamella. jSER: junctional SER, mito: mitochondrion. **(b)** 3D rendering of the JMC in (a) color-coded according to jSER width (grey to red) and T-tubule-jSER distance (blue range). **(c)** Quantification of T-tubule-jSER distance from 6 tomograms showing two populations of JMC distances. The peaks were fitted to Gaussian curves, and the peaks of the curves ± 2**σ** are indicated. **(d)** Quantification of jSER width. Thin jSER forms predominantly narrow JMC. The peaks were fitted to Gaussian curves, and the peaks of the curves ± 2**σ** are indicated for narrow and wide JMCs. **(e)** Close-up of a narrow JMC. White and black arrowheads mark rod-shaped JMC tethers and extracellular T-tubule densities, respectively. **(f)** Close-ups of wide JMCs. Prominent foot-like densities between jSER and T-tubule (white arrows) may correspond to RyRs. Black arrows mark jSER luminal densities. **(g)** Close-up of a tripartite JMC between a T-tubule, jSER and a mitochondrion. Abundant rod-shaped tethering densities are visible between T-tubule and jSER membranes (white arrowheads), as well as between jSER and mitochondrial membranes (grey arrowheads). Scale bars: 100 nm (a), 50 nm (e, f, g).

These data point to a model in which two distinct JMC domains coexist. On one hand, wide JMC may contain RyR2 channel clusters adjacent to a Ca2+ buffering jSER calsequestrin layer, thus specializing in trans-SER membrane Ca2+ handling. Adjacent narrow JMCs cannot spatially accommodate RyR2s and display numerous filamentous tethers, and thus may carry out different functions like non-vesicular lipid transfer. Interestingly, tripartite membrane contact sites between mitochondria, T-tubules and jSER were common only on thin JMC regions (Fig. 4g), hinting to complex multi-organelle communication at these junctions. Altogether, these first cryo-ET visualizations of native JMCs reveal novel morphological features and pave the way for future detailed molecular characterizations.

## Discussion

Our approach provides the first cryo-ET visualizations of unique cellular and tissue features that cannot be reproduced in culture. For example, we present here the first cryo-ET analyses of JMC in mature cardiomyocytes to date. On the other hand, *Drosophila* offers an extraordinary breadth of genetic tools to study basic biological mechanisms and model human disease (Bilder and Irvine, 2017, Ugur et al., 2016). Our method is also fully compatible with correlative cryo-light microscopy, but it also enables to target anatomical regions of interest directly by cryo-SEM imaging. Thus, we anticipate that the method presented here will allow addressing *in situ* the structural basis of a broad variety of biological phenomena. Since this method is limited to relatively small tissues, other techniques (Schaffer et al., 2019) remain valuable to study thicker tissue depths. On the other hand, our approach is not intrinsically limited to any specific sample type, and could also allow direct cryo-FIB/ET of, e.g., organoids, thin dissected mammalian tissues (Heuser and Tenkova, 2020) or even human biopsies, for example from the human heart (Brandenburg et al., 2022).

## Methods

### *Drosophila* tissues

#### Drosophila breeding, dissection and plunge freezing

Fly cultures were grown at 25 °C in standard fly medium. Assorted tissues from third instar larvae (L3 stage) of the wild type Canton-S strain were dissected during 2-5 min in PBS (Thermo Fisher) with or without 10 % glycerol after turning larvae inside out with the help of fine tip forceps. Dissections were performed under a Leica S6E stereomicroscope. Dissected tissues were placed on 200 mesh Molybdenum R2/1 EM grids (Quantifoil) immediately after dissection. The grids were subsequently mounted on a manual plunger, blotted from the back side using Whatman paper #1 (Sigma-Aldrich) and plunged into a 2:1 ethane:propane mixture cooled down by liquid nitrogen.

#### Cryo-FIB milling

To prepare thin electron transparent lamellae into the tissue, plunge-frozen grids were first mounted into autogrid frames (Thermo Fisher Scientific). The grids were then mounted into a dual-beam Quanta 3D FIB/SEM (FEI) using a custom-built transfer shuttle and a cryo-transfer system (PP3000T, Quorum). The samples were kept below -180 °C throughout FIB milling by the cryo-stage. To improve SEM imaging, a thin layer of pure metallic Pt was sputtered onto the sample under cryo conditions in the PP3000T transfer system to increase electrical conductivity. The following parameters were used: 10 mA sputtering current, 500 V between stage and sputtering target and 30 s of exposure at 4.5×10-2 mbar. To interpret and annotate the topographical anatomy of the tissue, overview maps of the EM grid were acquired by SEM at 10 kV at 100-250x magnification (object pixel size 1.1-0.4 µm) and by secondary electrons induced by the Ga+ focused ion beam at 30 kV and 338x magnification (object pixel size 0.7 µm). To protect the milling front of the lamellae, gaseous organic platinum was frozen on top of the grid using a gas injection system. To prevent bending of the lamella during the preparation, micro-expansion joints were milled left and right of the intended lamella preparation site (Wolff et al., 2019). 10-20 µm wide lamellae were prepared into the tissue with the ion beam at 30 kV at shallow angles (8-14°) in four consecutive steps: for the thicker tissue regions (e.g. VNC or skeletal muscle), the areas above and below the intended lamella were first removed with an ion beam current of 5 nA and 10 µm spacing. This step was not necessary for thinner tissues such as the peripheral nerves. Further rectangular patterns were defined above and below the intended lamella with 2 µm spacing for the rough milling step (ion beam current of 500-1000 pA), followed by fine milling with 800 nm spacing (100 pA) and a final polishing step down to the final lamella thickness of 100-200 nm (50 pA). To reach a uniform thickness, the lamella was tilted by ±0.5° and milled on each side separately with 50 pA current. The thickness of the lamella during the polishing step was assessed by SEM at 3-5 kV, 4.1 pA: vanishing of bright (charged) areas of the lamella indicated a thickness of <300 nm at 5 kV or <200 nm at 3 kV. Biological structures inside the lamella at the surface were imaged at each step by SEM at 2.5 kV, and4.1 pA in integration mode (64x). To reduce lamella charging during phase plate cryo-ET data acquisition, a thin layer of pure metallic Pt was sputtered onto the lamella under cryo conditions in the PP3000T transfer system with the following parameters: 5 mA sputtering current, 500 V between stage and sputtering target and 10 s of exposure at 4.5×10-2 mbar.

#### Cryo-TEM imaging and diffraction

Cryo-FIB lamellae were imaged using a Titan Krios cryo-TEM (FEI) equipped with a field emission gun operated at 300 kV, a Volta phase (Danev et al., 2014), a post-column energy filter (Gatan) operated at zero-loss and a K2 Summit direct electron detector (Gatan). Low magnification projections of lamellae were recorded (11.500 x, object pixel size 1.312 nm for Fig. 2m and Extended Data Fig. 3e; 4.800 x, object pixel size 3.064 nm for all other lamella overviews) using the montage option in SerialEM (Mastronarde, 2005) (RRID:SCR_017293) with 15% overlap and -20 µm defocus. Phase plate alignment and operation was carried out as described (Fukuda et al., 2015). Upon phase plate conditioning, high magnification (33,000x, object pixel size 0.421 nm) tilt series were recorded at locations of interest using the SerialEM low dose acquisition scheme with a tilt increment of 2°, typically spanning an angular range from -60+ξ° to +60+ξ°, where ξ is the pre-tilt of the lamella (∼+13°). Target defocus was set to -0.5 µm. The K2 camera was operated in dose fractionation mode recording frames every 0.2 s. For each tilt series, a new spot on the phase plate was selected. The total dose was limited to 100-150 e-/Å2. Diffraction patterns were acquired of the indicated regions with the same beam settings as for tomography, a selected area of 7 µm2, a camera length of 1.95 m and a total dose of 10 e-/Å2.

#### Tomogram reconstruction and segmentation

Raw K2 camera frames were aligned using MotionCor2 (Zheng et al., 2017) (RRID:SCR_016499). The resulting tilt series were aligned using patch tracking in version 4.7 of the IMOD package (Kremer et al., 1996) (RRID:SCR_003297) and reconstructed by weighted back projection. For tomographic reconstruction, a radial filter with the following values was applied: cut-off: 0.35, fall-off: 0.05. Membranes were first automatically segmented using the TomoSegMemTV package (Martinez-Sanchez et al., 2014) and refined manually and visualized using Amira (FEI Visualization Sciences Group; RRID:SCR_014305). Microtubules and the sarcomeric cytoskeleton were automatically detected using the XTracing Module in Amira (Rigort et al., 2012b).

### Mouse cardiomyocytes

#### Cardiomyocyte isolation and vitrification

Living adult wild-type C57Bl6N mouse ventricular cardiomyocytes were isolated from Langendorff-perfused hearts by collagenase digestion using established workflows tuned to preserve subcellular structural integrity (Wagner et al., 2012, Wagner et al., 2014). Briefly, the heart was Langendorff perfused with Ca2+-free oxygenated Krebs solution (perfusion buffer; 120.4 mM NaCl, 14.7 mM KCl, 0.6 mM KH2PO4, 0.6 mM Na2HPO4, 1.2 mM MgSO4, 10 mM HEPES, 4.6 mM NaHCO3, 30 mM taurin, 10 mM 2,3-butanedione-monoxime, 5.5 mM glucose, pH 7.4) at 42°C for 2 minutes. The perfusion buffer was then switched to collagenase type II containing solution for 12 minutes (perfusion buffer supplemented with 2 mg/ml collagenase and 40 μM CaCl2). Following digestion, the atria were carefully excised and discarded. The ventricles were dissected into 1 mm3 pieces in the collagenase-containing buffer; digestion was stopped using perfusion buffer supplemented with 10% bovine calf serum and 12.5 μM CaCl2 (stop buffer). The cells were washed twice (once with stop buffer and once with perfusion buffer), inspected on aby light microscope microscopy before plating on EM grids.

Prior to vitrification, 200 mesh R2/1 Au EM grids (Quantifoil) were coated with laminin (1:10 dilution) for 30 minutes, followed by a brief wash in PBS. Immediately after CM preparation, the cells were gently resuspended, were applied to the coated grids and left to sediment for 15 minutes. The cells were inspected on a light microscope for good cell isolation quality and grid-attachment quality. To remove dead and non-sedimented cells, grids were washed by applying a drop of perfusion buffer and gently pipetting up and down. Finally, a drop of perfusion buffer supplemented with 10% glycerol was applied to the grids for no longer than one minute, the grids were mounted on a manual plunging device, blotted from the back, and vitrified by plunging into a 2:1 ethane:propane mixture cooled by liquid nitrogen.

#### Cryo-FIB milling

Plunge-frozen grids with CMs were clipped onto cryo-FIB autogrids (Thermo Fisher Scientific) and then mounted onto a specimen shuttle for insertion into an Aquilos 2 Cryo-FIB (Thermo Fisher Scientific). The shuttle was transferred to the microscope via a cryo-transfer rod (Thermo Fisher Scientific). The samples were kept under -180 °C throughout FIB milling by the cryo-stage. A layer of metallic Pt was sputtered onto the sample for 15 s with a current of 30 mA under cryo-conditions, followed by application of organometallic Pt for 45 s using the gas injection system to protect the front of the lamellae during milling. For rough milling, rectangular patterns were defined above and below the intended lamella with 3 µm spacing (ion beam current of 500-1000 pA). Then, using three line milling patterns (30 s at a current of 100 pA), a notch was milled into the flank of the rough lamella to allow micro-expansion and prevent bending during fine milling (Kelley et al., 2022). Further milling was carried out by gradually reducing the spacing between the milling areas and simultaneously reducing beam current. Final polishing of lamellae was performed with a beam current of 30 pA to a thickness of 100-200 nm. To counteract charging induced motion, a layer of metallic Pt was sputtered on to the fine-milled lamellae (3 s at a current of 30 mA).

#### Tilt-series data collection and processing

Tilt-series of CM lamellae were collected on a 300 kV Krios G4 cryo-TEM equipped with a Falcon 4 detector and a Selectris energy filter (Thermo Fisher Scientific), using a slit width of 15 eV. Montages of lamellae were recorded at a magnification of 4,800x using the inbuilt SerialEM function (Mastronarde, 2005). High-magnification tilt-series were recorded with the PACEtomo plugin for SerialEM (Eisenstein et al., 2022) at 42,000x, corresponding to a 2.94 Å pixel size. Starting from the pre-tilt of the lamellae (∼13°), tilt series were collected using a dose-symmetric tilt scheme with a step size of 3° for a total of 37 tilt movies and a combined dose of 120-150 e-/Å2. Motion correction and CTF estimation were carried in WARP v.1.0.9 (Tegunov and Cramer, 2019). Tilt series were aligned using bead tracking of Pt components on the lamellae surface and reconstructed in version 4.7 of the IMOD package by weighted back projection (Kremer et al., 1996). We utilized MaskTomRec (Fernandez et al., 2016) to computationally remove the layer of sputtered Pt on the lamellae surfaces and CryoCARE (Buchholz et al., 2019) for denoising of tomograms. For visualization, tomograms were deconvoluted using a Wiener-like filter (Tegunov and Cramer, 2019).

#### Tomogram analysis

Membranes from 6 tomograms were initially automatically segmented using the TomoSegMemTV package (Martinez-Sanchez et al., 2014) followed by manually selecting and cleaning junctional sarcoplasmic reticulum and T-tubule membranes with Amira (Thermo Fisher Scientific). The membrane segmentations were converted into meshes and distance measurements between reconstructed mesh triangles were carried out using a previously established surface morphometrics pipeline (Barad et al., 2023, Salfer et al., 2020). Per-triangle measurements of T-tubule-jSER distances were weighted by surface area and plotted as histograms using matplotlib (Harris et al., 2020, Hunter, 2007). jSER thicknesses were measured as distances between proximal and distal jSER membranes and plotted separately for narrow JMC regions and wide JMC regions. The peaks were fitted with Gaussian curves: for the T-tubule-jSER distance distribution the fitting was done with the sum of two Gaussian curves and for the narrow and wide JMC each with a single Gaussian curve (adj. R2 > 0.9). The peak maximum and the value of the standard deviation were derived from the fitting results.

#### Lamellae stitching and clearing (LisC filtering)

Low magnification images of lamellae were stitched using the Image Composite Editor (Microsoft Research) to produce complete overviews. Image tiles were imported using the option “Structured Panorama”. The camera motion was defined to “Planar motion”, the image order “Zigzag”, the angular range to “Less than 360°”, the overlap according to the parameter set in the montage menu of SerialEM and the search radius 5% more than the overlap. After the stitching procedure, the lamella was rotated so that the curtains were vertically aligned, and the overviews were cropped and saved.

The LisC algorithm was programmed as macro for Fiji (Schindelin et al., 2012) (RRID:SCR_002285) and is publicly available (https://github.com/FJBauerlein/LisC_Algorithm). The input for the macro are the stitched lamella overview image and its pixel size. First, the image is high-pass filtered to remove the low frequency brightness modulations caused by long-range thickness variations. For this, the Fiji function “Bandpass filter” is used with the cut-off values of 1 pixel and 5 μm/pixel size. Then, masks for regions of extreme contrast (e.g. ice crystal contamination and vacuum) are created to avoid ringing artifacts caused by filtering in Fourier space. For that, thresholds for the contamination (default: mean - 1.5xSD) and vacuum (default: mean + 1.5xSD) regions are set. The masks are then applied on the original image and the pixel values of the masked regions are substituted by a local gray value average obtained by a Gaussian blur filter with a sigma of 5 μm divided by the pixel size. Then, the line artifacts caused by curtaining are removed by a line filter, where the line in Fourier space is placed perpendicular to the line artifacts in real space. This is applied by the Fiji function “Suppress stripes” with a tolerance of direction of 95%. The Fourier filtering also causes gray values changes of the contamination and vacuum regions defined by the mask. Therefore, the pixel values in those regions are set to 0 (contamination) and 256 (vacuum), respectively.

## Data availability

The tomograms shown in Fig. 2 are available at EMDB with the following accession codes: EMD-12726 (Fig. 2c, d), EMD-12727 (Fig. 2f, g), EMD-12728 (Fig. 2j, k) and EMD-12729 (Fig. 2n, o).

## Code availability

The LisC algorithm is available at:

https://github.com/FJBauerlein/LisC_Algorithm.

## Acknowledgments

We thank Nicolas Gompel and Bettina Mühling for providing *Drosophila* cultures. We thank Tobias Kohl for support with the protocol optimization for the cardiomyocyte plating on EM grids. We thank Wolfgang Baumeister and Jürgen Plitzko for providing access to electron microscopy infrastructure at the Max Planck Institute of Biochemistry and stimulating discussions. We thank Philipp Erdmann, Miroslava Schaffer and Tat Cheng for electron microscopy support. S.E.L. and R.F.-B. acknowledge funding from the Deutsche Forschungsgemeinschaft (DFG, German Research Foundation) through Germany’s Excellence Strategy - EXC 2067/1-390729940 and SFB1190/P03 (S.E.L) and P22 (R.F.-B). Cryo-ET instrumentation at the University of Göttingen was jointly funded by the DFG Major Research Instrumentation program (448415290) and the Ministry of Science and Culture of the State of Lower Saxony. J.C.P.-P. acknowledges funding from Natural Science Foundation of China (NSFC) grants 91854207 and 32150710524 and the Spanish Ministry for Science and Innovation through grant PID2021-122119NB-I00 and the “Severo Ochoa” Program for Centers of Excellence (CEX2021-001165-S).

## Author contributions

F.J.B.B., S.E.L., J.C.P.-P. and R.F.-B. planned research. J.C.P.-P. performed *Drosophila* dissections. D.E.C. prepared cardiomyocyte samples. F.J.B.B. and M.R. vitrified samples and performed cryo-FIB and cryo-ET. F.J.B.B. and M.R. analyzed the data. J.C.P.-P. and R.F.-B. supervised electron microscopy experiments and data analysis. F.J.B.B., S.E.L., J.C.P.-P. and R.F.-B. wrote the manuscript with contributions from all authors.

## Additional information

Supplementary Information is available for this paper.

Correspondence and requests for materials should be addressed to

F.J.B.B.,

S.E.L.

J.C.P.-P. or

R.F.-B.

---

**Extended Data Fig. 1:**
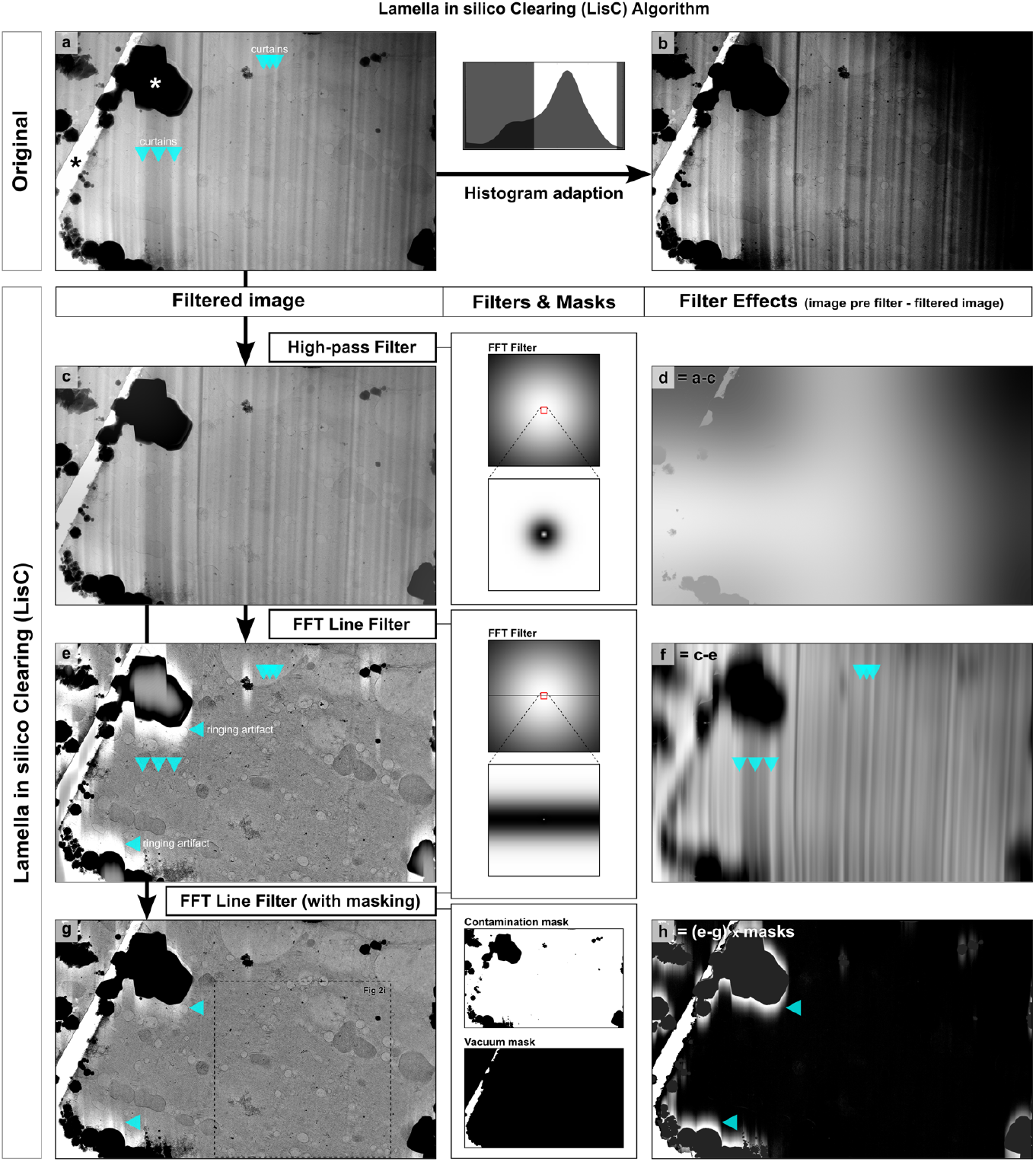
Lamella in-silico Clearing (LisC) algorithm. The left column shows images before and after the different filtering steps. The middle column illustrates the filters and masks applied. The right column shows the effect of the filter, i.e. the image pre-filter minus the filtered image. **(a)** Original overview micrograph of a lamella showing typical cryo-FIB milling imperfections such as ‘curtains’ (blue downwards arrows), i.e. local stripe-like variations in thickness, as well as long-range thickness differences over the whole lamella. These medium- and low-frequency thickness variations cause strong contrast alterations and make it difficult to identify biological structures in the lamella. A white asterisk marks ice contamination on top of the lamella, and a black asterisk marks the vacuum region, which can be seen through a crack in the lamella. **(b)** These artifacts become more pronounced if contrast is increased. **(c)** A high-pass filter removes the low frequency brightness modulation caused by the long-range thickness variations. The filter is shown in Fourier space in the box ‘High-pass Filter’. A red box marks the region magnified below. **(d)** Difference of the high-pass filtered image and the original: (d) = (a) – (c). **(e)** The anisotropic, unidirectional curtains are efficiently removed (blue down arrows) by a line mask in Fourier space (see box ‘FFT Line Filter’) perpendicular to the direction of the curtains. A red box marks the region magnified below. The very strong contrast gradients between the lamella and ice crystal contamination or the vacuum regions marked by asterisks in (a) cause ringing artifacts (blue lateral arrows). **(f)** Difference of the FFT line-filtered image and the high-pass filtered image: (f) = (c) – (e). The filter efficiently removes curtains without affecting biological structures in the lamella. **(g)** The ringing artifacts shown in (e) can be strongly reduced by applying masks (see box ‘FFT Line Filter (with masking)’) to set the pixel values of the contamination and vacuum regions to a local gray value average before FFT line filtering. Lastly, the pixel values in the masked regions are set to 0 (contamination) and 256 (vacuum), respectively. The structural features of the tissue can be clearly discerned, as annotated in the area of this lamella shown in Fig. 2i (white box). **(h)** Difference of the masked vs. the unmasked FFT line-filtered image. The pixel values in the masked regions are set to 0 (contamination) and 256 (vacuum): (h) = [(e) – (g)] x masks. The LisC algorithm is publicly available (https://github.com/FJBauerlein/LisC_Algorithm) as a macro for Fiji. See also Supplementary Video 2.

**Extended Data Fig. 2:**
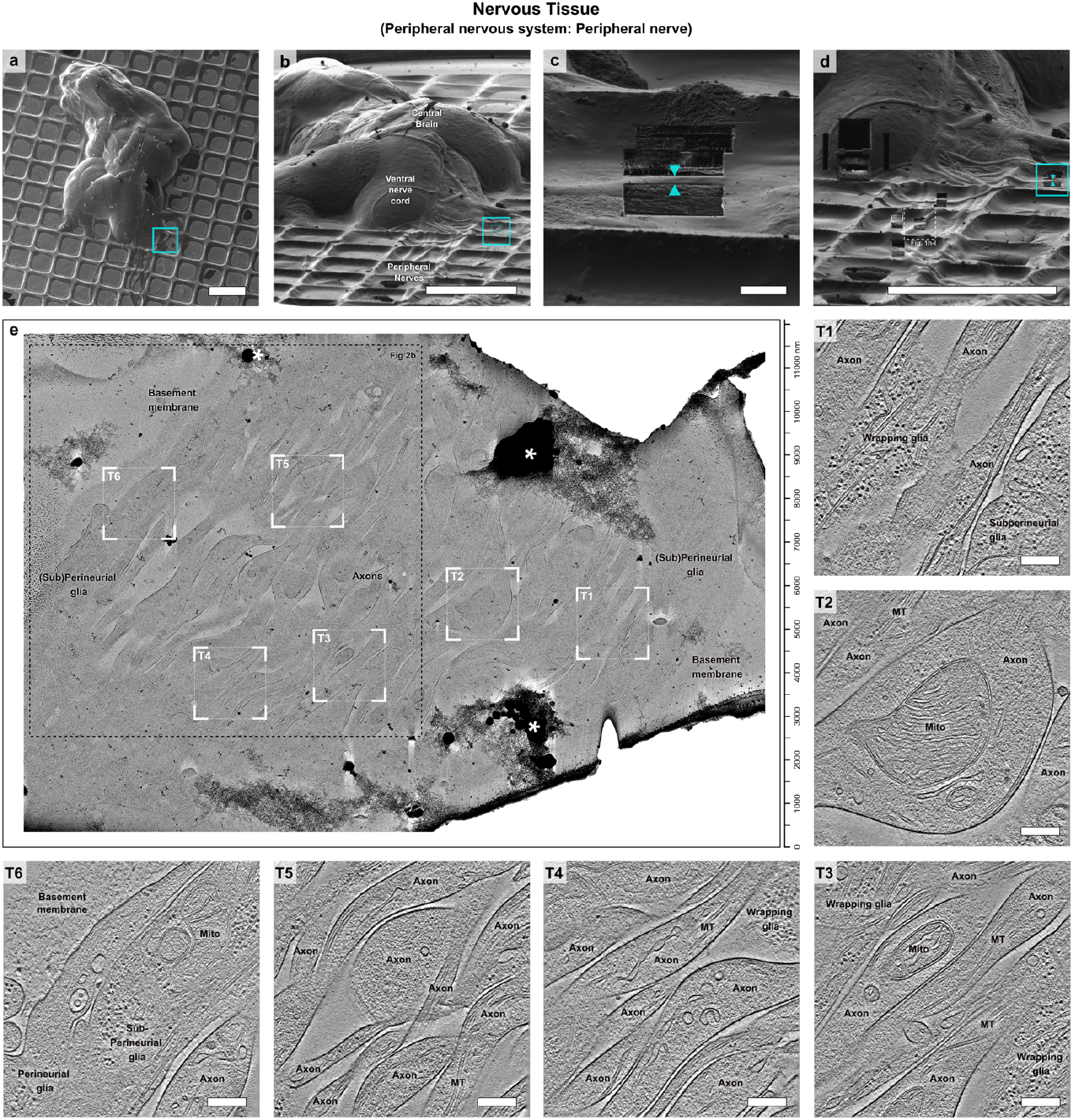
Cryo-FIB lamella preparation of nervous tissue for cryo-ET imaging. **(a)** Top-view cryo-SEM image of the *Drosophila* cephalic complex shown in Fig. 1g and Fig. 2a. **(b)** Cryo-FIB-induced secondary electron side view of (a). **(c)** Preparation of the lamella shown in Fig. 2b on a peripheral nerve at the location marked by a blue rectangle in (b). The lamella is indicated by blue arrows. **(d)** Cryo-FIB overview of the tissue with the final ∼150 nm thick lamella (blue box). **(e)** Cryo-TEM overview of the final lamella, visualizing parallel axons, sub-perineurial and perineurial glia, delimited by the basement membrane. The image was filtered using the LisC algorithm. The area of the lamella shown in Fig. 2b is indicated by a black box. Tomograms were acquired in the regions marked by white boxes. **(T1-T6)** 1.7 nm-thick slices of the tomograms recorded in the regions marked in (e). Mito: mitochondrion; MT: microtubule. T6 is also shown in Fig. 2c, d and Supplementary Video 3 with its corresponding 3D rendering. Scale bars: 200 μm (a, b, d), 10 μm (c), 250 nm (T1-T6).

**Extended Data Fig. 3:**
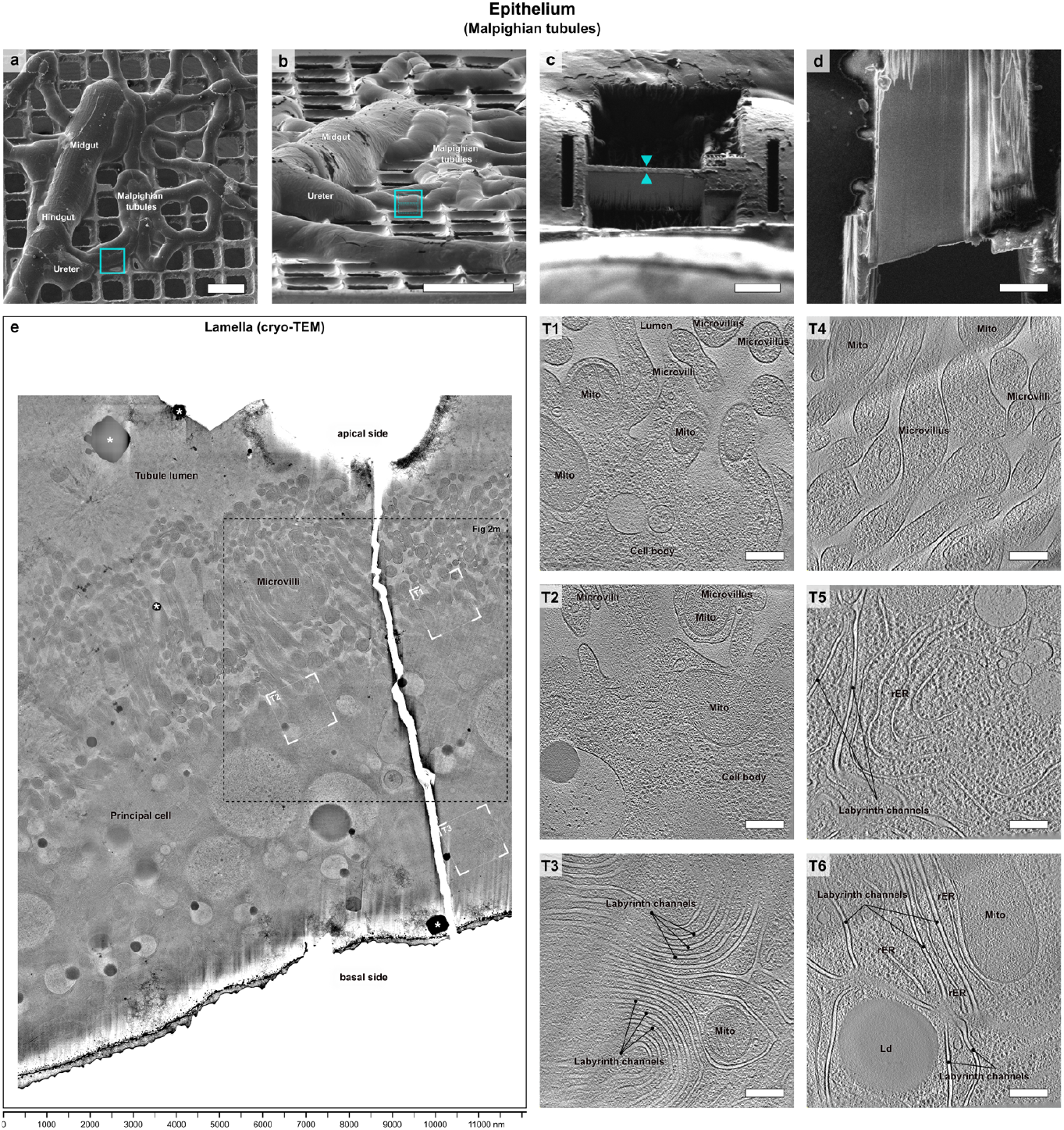
Cryo-FIB lamella preparation of epithelial tissue for cryo-ET imaging. **(a)** Top-view cryo-SEM image of the *Drosophila* Malpighian tubules shown in Fig. 2l. **(b)** Cryo-FIB-induced secondary electron side view of (a). **(c)** Preparation of the lamella shown in Fig. 2m on a Malpighian tubule at the location marked by a blue rectangle in (b). The lamella is indicated by blue arrows. **(d)** Cryo-SEM image of the final lamella. **(e)** Cryo-TEM overview of the final lamella, visualizing a principal cell from the basal to the apical pole with multiple microvilli emerging into the tubule lumen. The image was filtered using the LisC algorithm. The area of the lamella shown in Fig. 2m is indicated by a black box. Tomograms were acquired in the regions marked by white boxes. **(T1-T3)** 1.7 nm-thick slices of the tomograms recorded in the regions marked in (e). T1 is also shown in Fig. 2n, o and Supplementary Video 6 with its corresponding 3D rendering. **(T4-T6)** 1.7 nm-thick slices of the tomograms recorded on another lamella. Ld: lipid droplet; Mito: mitochondrion; rER: rough endoplasmic reticulum. Scale bars: 200 μm (a, b), 10 μm (c, d), 250 nm (T1-T6).

